# An acetylation-mediated chromatin switch governs H3K4 methylation read-write capability

**DOI:** 10.1101/2022.02.28.482307

**Authors:** Kanishk Jain, Matthew R. Marunde, Jonathan M. Burg, Susan L. Gloor, Faith M. Joseph, Karl F. Poncha, Zachary B. Gillespie, Keli L. Rodriguez, Irina K. Popova, Nathan W. Hall, Anup Vaidya, Sarah A. Howard, Hailey F. Taylor, Laylo Mukhsinova, Ugochi C. Onuoha, Emily F. Patteson, Spencer W. Cooke, Bethany C. Taylor, Ellen N. Weinzapfel, Marcus A. Cheek, Matthew J. Meiners, Geoffrey C. Fox, Kevin E. W. Namitz, Martis W. Cowles, Krzysztof Krajewski, Zu-Wen Sun, Michael S. Cosgrove, Nicolas L. Young, Michael-Christopher Keogh, Brian D. Strahl

## Abstract

In nucleosomes, histone N-terminal tails exist in dynamic equilibrium between free/accessible and collapsed/DNA-bound states. The latter state is expected to impact histone N-termini availability to the epigenetic machinery. Notably, H3 tail acetylation (*e.g.*, K9ac, K14ac, K18ac) is linked to increased H3K4me3 engagement by the BPTF PHD finger, but it is unknown if this mechanism has broader extension. Here we show that H3 tail acetylation promotes nucleosomal accessibility to other H3K4 methyl readers, and importantly, extends to H3K4 writers, notably methyltransferase MLL1. This regulation is not observed on peptide substrates yet occurs on the *cis* H3 tail, as determined with fully-defined heterotypic nucleosomes. *In vivo*, H3 tail acetylation is directly and dynamically coupled with *cis* H3K4 methylation levels. Together, these observations reveal an acetylation ‘chromatin switch’ on the H3 tail that modulates read-write accessibility in nucleosomes and resolve the long-standing question of why H3K4me3 levels are coupled with H3 acetylation.

## INTRODUCTION

In the epigenetic landscape, histone proteins are often variably chemically modified by “writer” enzymes (Jenuwein and Allis, 2001; Strahl and Allis, 2000). Writer-installed post-translational modifications (PTMs) can then be recognized by “reader” proteins and/or removed by “eraser” enzymes. This interplay of PTMs comprises the “histone code,” and has a central function in regulating chromatin organization and activity. For example, methylated/acylated lysine or methylated arginine residues of histones can recruit transcription factors to activate or repress transcription (Strahl and Allis, 2000; Su and Denu, 2016); mitotically phosphorylated serine/threonine residues can regulate reader binding established at earlier stages of the cell cycle (Rossetto et al., 2012); or ubiquitinated lysine can impact the maintenance of DNA methylation (Vaughan et al., 2021). As the complex language of histone PTMs is dissected, it has become clear that multivalent interactions with reader proteins can influence chromatin structure and DNA accessibility, thereby regulating gene transcription and other DNA-templated events (Su and Denu, 2016; Taylor and Young, 2021; Young et al., 2010). In this manner, combinatorial PTMs can more effectively engage different chromatin-binding modules, promoting distinct outcomes versus either PTM alone.

The bulk of chromatin PTM research has employed histone peptides, even though histones exist *in vivo* in a heteromeric complex with DNA (*i.e.*, the nucleosome). Recent work, however, is making it increasingly clear that studying histone PTM engagement in the nucleosome context provides a more accurate understanding of the histone code. Particularly, the highly charged histone tails interact directly with nucleosomal DNA, restricting access for PTM recognition by reader proteins (Ghoneim et al., 2021; Marunde et al., 2022a; Morrison et al., 2018). Studies with BPTF PHD suggest acetylation releases the H3 N-terminal tail from the nucleosome surface, such that H3K4me3 becomes more readily engaged by the PHD finger (Marunde et al., 2022a; Morrison et al., 2018).

Considerable research effort has focused on dissecting the direct (and multivalent) engagement of chromatin via histone PTM-reader protein interactions. However, less appreciated are any indirect effects of PTMs on histone tail accessibility/nucleosome dynamics (*e.g.,* via charge neutralization). In this report, we demonstrate enhanced nucleosome binding by a range of H3K4 readers when the histone tail is concomitantly acetylated (one or more of K9ac, K14ac, and K18ac). Furthermore, from *in vitro* enzymatic assays, we found that neighboring acetylation of the *cis* H3 tail is a prerequisite switch that enables the MLL1 complex (MLL1C: MLL1 SET domain, WDR5, RbBP5, Ash2L and DPY30) (Rao and Dou, 2015) to robustly methylate H3K4. Consistent with this observation, mass spectrometric proteomic analyses of mammalian cells in a timed response to sodium butyrate (a broad-spectrum lysine deacetylase (KDAC) inhibitor) revealed a tight correlation of H3K4 methylation with *cis* acetylation. Our findings define a critical aspect of chromatin regulation: *i.e.,* PTM crosstalk through acetylation-mediated tail accessibility. The findings also provide a molecular basis for the long-standing connection between H3K4 methylation and H3 acetylation in multiple eukaryotes (Garcia et al., 2007; Nightingale et al., 2007; Taverna et al., 2007), and resolve the directionality of these correlations: *cis* hyperacetylation of the H3 tail precedes, and is largely a prerequisite for, H3K4 methylation. Thus, the establishment of sites of H3K4me3 and activation of transcription occur by a sequence of modifications of the same histone molecule.

## RESULTS

### PHD finger readers show narrowed selectivity for histone tail PTMs on mononucleosomes versus peptides

How histone readers engage nucleosomes is an extensively researched area of chromatin biology. Most investigators characterize reader binding with PTM-defined histone peptides, although the domains often display a refined preference to similarly modified nucleosomes (Marunde et al., 2022b, 2022a; Morgan et al., 2021; Morrison et al., 2018). To further assess this potential, we used the dCypher^®^ approach (Jain et al., 2020; Marunde et al., 2022b; Morgan et al., 2021; Weinberg et al., 2021) to measure the interactions of three PHD readers (from KDM7A, DIDO1, and MLL5: the queries) with PTM-defined peptides and nucleosomes (the potential targets). As might be expected (Jain et al., 2020), each GST-PHD fusion showed a preference for H3K4-methylated peptides and, particularly, to higher methyl states (*i.e.*, KDM7A bound me2/me3, DIDO1 bound me1/me2/me3, and MLL1 bound me1/me2/me3; **Figure 1A**). In contrast, each GST-PHD reader was restricted to H3K4me3 over the lower methyl states on nucleosomes (**Figure 1B**) and displayed weaker relative binding (EC_50_^rel^: calculated as in **Methods**) to this PTM (**Figure 1 - figure supplement 1C-D**). On further examination, we observed no impact of co-incident acetylation on H3K4me3 binding in the peptide context (**Figure 1A**: compare H3K4me3 to H3K4me3K9acK14acK18ac [hereafter H3K4me3tri^ac^]). In stark contrast, the binding of each GST-PHD reader to nucleosomal H3K4me3 was dramatically enhanced (∼10-15-fold) by co-incident acetylation (*i.e.*, H3K4me3tri^ac^; **Figure 1B** and **Figure 1 - figure supplement 1A-B**). Additionally, there was no significant reader domain interaction with either nucleosomes lacking H3K4 methylation or 147X601 DNA (nucleosomal DNA) alone (**Figure 1—figure supplement 1B**). A similar observation has also been made for the BPTF PHD domain (Marunde et al., 2022a; Morrison et al., 2018), suggesting potential for a general mechanism. This led us to consider the possibility that histone tail lysine acetylation (Kac) might function beyond the recruitment of readers, and perhaps also impact H3K4 writers.

**Figure 1.**
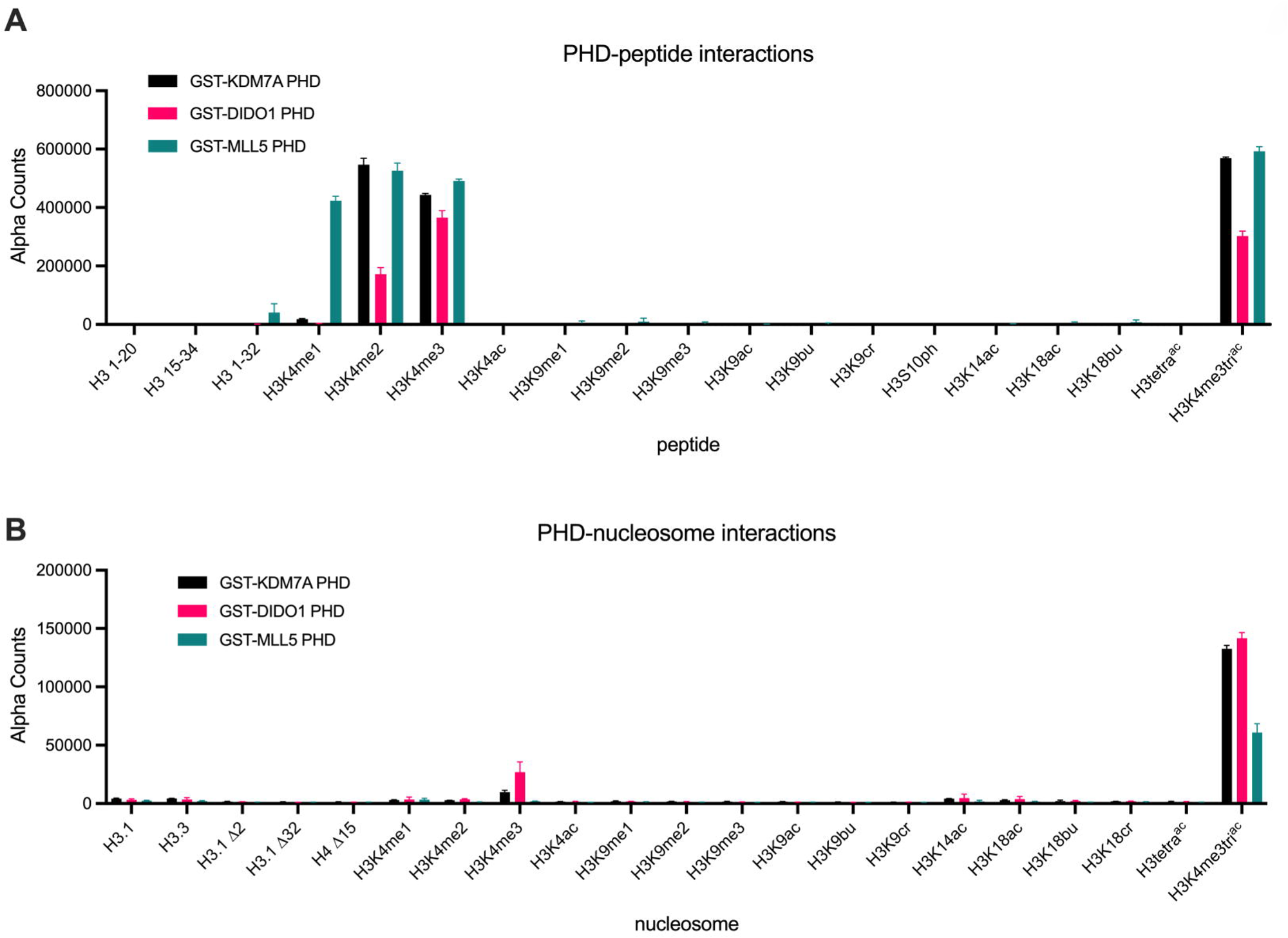
PHD finger reader domains show restricted binding on PTM-defined peptides vs. nucleosomes. *dCypher* assay alpha counts for interaction of GST-PHD queries (9.5 nM KDM7A (Uniprot #Q6ZMT4; residues 1-100); 2.4 nM DIDO1 (Uniprot #Q9BTC0; residues 250-340); 18 nM MLL5 (Uniprot #Q8IZD2; residues 100-180)) with PTM-defined peptides (**A**) *vs.* nucleosomes (**B**) (the potential targets). All error bars represent the range of two replicates. Key: H3.1 Δ2, H3.1 Δ32 and H4 Δ15 are nucleosomes assembled with histones lacking the indicated N-terminal residues of H3.1 or H4. All data was plotted using GraphPad Prism 9.0. Of note, each reader query showed minimal interaction free DNA (147bp or 199bp: **Figure 1 – figure supplement 1B)**.

### H3 N-terminal acetylation enhances MLL1C mediated methylation of nucleosomal H3K4

To investigate if acetylation might enable a more catalytically accessible H3 N-terminus, we performed enzymatic assays with the MLL1 core complex (MLL1C; responsible for H3K4me3) (Rao and Dou, 2015; Sha et al., 2020) and defined nucleosome substrates ± accompanying acetylation (H3K9acK14acK18ac; hereafter H3tri^ac^) (see **Methods**). In an endpoint assay at constant enzyme and substrate concentrations (and [*methyl*-^3^H]-SAM donor), we observed a significant increase in net methylation when the H3 tail was also acetylated (**Figure 2A**). As expected, methylation by MLL1C sequentially decreased towards H3K4 mono-, di-, and tri-methylated nucleosomal substrates, being undetectable on H3K4me3 (which also confirmed MLL1C targeting of this specific residue). Despite this, methyl group incorporation to each H3K4 methyl state substrate was consistently enhanced by accompanying H3tri^ac^ (**Figure 2A**). Of note, we also tested the viability of methyllysine analogs (MLAs) (Simon and Shokat, 2012) ± H3tri^ac^ as MLL1C substrates and observed no activity, indicating their unsuitability for such studies (**Figure 2 - figure supplement 1D-E**).

**Figure 2.**
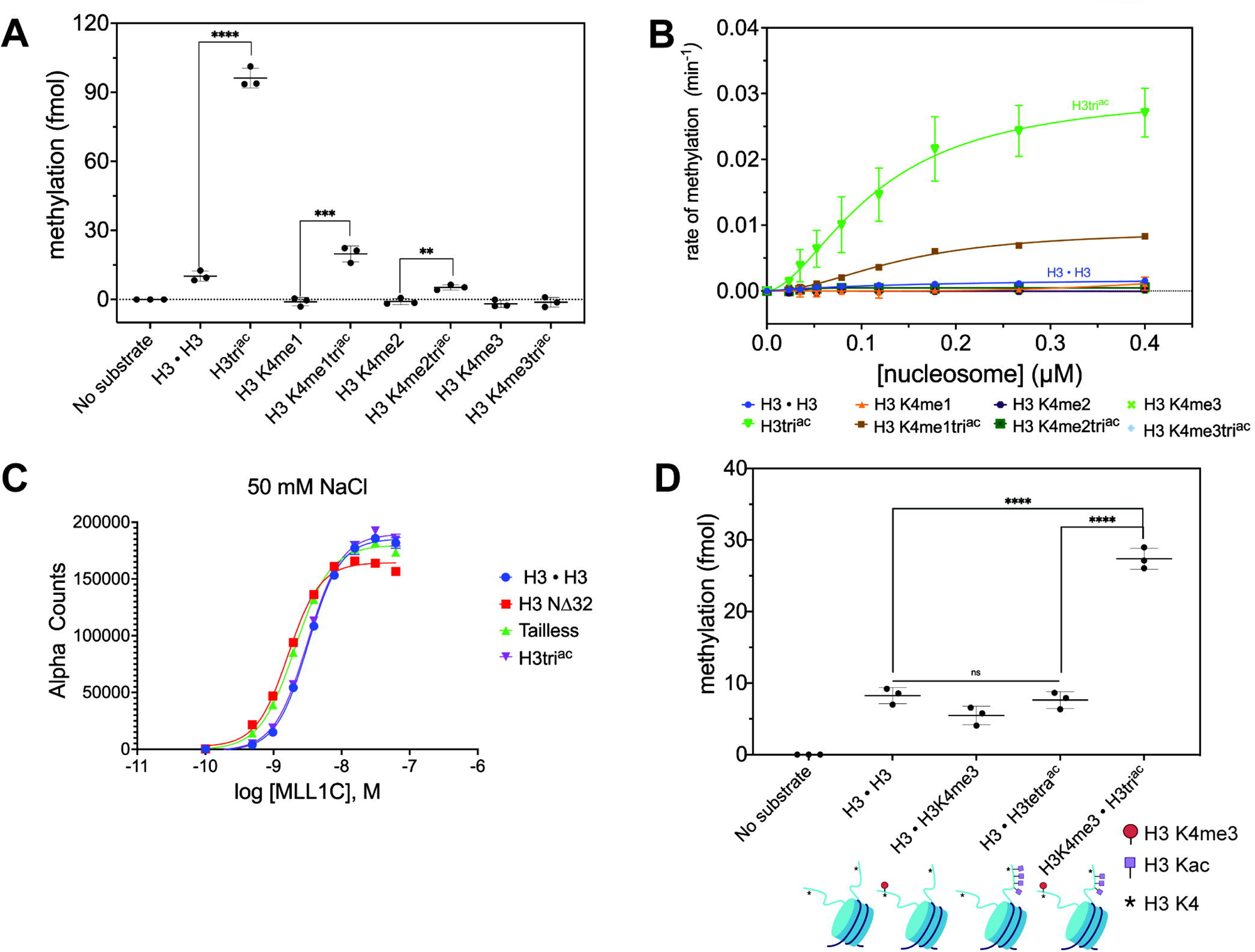
MLL1C methylation activity on nucleosomal H3K4 is significantly enhanced by co-incident H3tri^ac^. **A)** Endpoint methylation assays of H3K4me0 (H3 • H3)-me1-me2-me3 nucleosomes and their cognate H3K9ac14ac18ac (H3tri^ac^) partners (all 100 nM) with MLL1 Complex (MLL1C; 4nM). Reactions performed in triplicate with error bars as S.E.M. *p*-values were determined using a two-tailed t test: **** = <0.0001, *** = 0.0008, ** = 0.0038. **B)** MLL1C (4 nM) methylation activity on H3K4me0-me1-me2-me3 nucleosomes and their cognate H3triac partners (all substrates: 1.5-fold serial dilution, 0-0.4 μM). Reactions performed in triplicate with error bars as S.E.M. (see also **Figure 2 - figure supplement 1**). **C)** MLL1C does not differentially associate with PTM-defined nucleosome substrates under study conditions. dCypher binding curves of 6HIS-tagged MLL1C (concentrations noted) with PTM-defined nucleosomes (20 nM). Error bars represent the range of two replicates. **D)** MLL1C-mediated methylation is enhanced on cis but not trans acetylated nucleosomal H3 tails. Endpoint methylation assays of MLL1C (4nM) with homotypic [H3 • H3] *vs.* heterotypic (e.g. [H3 • H3tetra^ac^]; see **Methods**) nucleosome substrates (all 100 nM). Reactions performed in triplicate with error bars as S.E.M. *p*-values were determined using a two-tailed t test: **** = <0.0001. Key: H3tri^ac^ = H3K9acK14acK18^ac^. H3tetra^ac^ = H3K4acK9acK14acK18ac.

We next measured the steady-state methylation kinetics of MLL1C towards nucleosomes with each H3K4 methyl state ± H3tri^ac^, and again observed that acetylation increased methyltransferase activity (**Figure 2B**). Using an extra sum-of-squares F-test, methylation for the H3tri^ac^ nucleosomes was indicative of positive cooperativity because of better fit (*p* = 0.0216) to the Hill equation (Weiss, 1997) (compare **Figure 2B** and **Figure 2 - figure supplement 1C**; **Table 1** and **Supplementary File 1**). Because of the low level of enzymatic activity towards the unacetylated nucleosomes we could not make a statistically significant comparison between the Hill and Michaelis-Menten fits. There have been limited studies of MLL1C activity on nucleosomes (Park et al., 2019; Patel et al., 2011; Xue et al., 2019), so an overlooked potential allostery is understandable given the many possible interactions between this enzyme complex and substrate (Lee et al., 2021; Park et al., 2019).

**TABLE 1:** Steady-state Hill kinetic parameters

| Substrate | $K_{0.5}$ ($\mu\text{M}$) | $k_{\text{cat}}$ ($\text{min}^{-1}$) | $h$ (Hill coefficient) | $R^2$ |
| --- | --- | --- | --- | --- |
| H3 • H3 | $0.13 \pm 0.06$ | $0.0018 \pm 0.0004$ | $1.213 \pm 0.3430$ | 0.8810 |
| H3<br>K9ac/14ac/18ac | $0.12 \pm 0.02$ | $0.0304 \pm 0.0036$ | $1.741 \pm 0.3528$ | 0.9228 |
| H3K4me1 | n.d.<sup>a</sup> | n.d.<sup>a</sup> | n.d.<sup>a</sup> | 0.3186 |
| H3K4me1<br>K9ac/14ac/18ac | $0.14 \pm 0.008$ | $0.0092 \pm 0.0004$ | $2.032 \pm 0.1333$ | 0.9920 |
| H3K4me2 | n.d.<sup>a</sup> | n.d.<sup>a</sup> | n.d.<sup>a</sup> | -0.1746 |
| H3K4me2<br>K9ac/14ac/18ac | $0.042 \pm 0.005$ | $0.0005 \pm 0.00004$ | $5.349 \pm 2.682$ | 0.8155 |
| H3K4me3 | n.d.<sup>a</sup> | n.d.<sup>a</sup> | n.d.<sup>a</sup> | n.d.<sup>a</sup> |
| H3K4me3<br>K9ac/14ac/18ac | n.d.<sup>a</sup> | n.d.<sup>a</sup> | n.d.<sup>a</sup> | n.d.<sup>a</sup> |

<sup>a</sup> = methylation signal was indistinguishable from background: kinetic parameters could not be determined.

On examining enzymatic parameters in detail, we noted that although overall kcat was ∼17-fold greater for homotypic H3tri^ac^ (over unmodified, H3 • H3) nucleosomes, the K_0.5_ values (substrate concentration at half-maximal velocity/half-saturation for an allosterically regulated enzyme) were indistinguishable (**Table 1**). Therefore, although MLL1C catalytic efficiency toward H3tri^ac^ nucleosomes was enhanced by an increase in k_cat_, this catalytic efficiency was not driven by K_0.5_, which suggested no increase in relative binding affinity. To further examine this, we returned to the dCypher approach to examine potential binding between MLL1C query and a selection of nucleosome targets: H3 • H3, H3NΔ32 (lacking the first 32 residues of H3), Tailless (trypsin-digested nucleosomes to remove N- and C-terminal histone tails), and H3tri^ac^. At 50 mM NaCl, we observed no compelling difference in MLL1C binding to any of these targets (**Figure 2C** and **Figure 2 - figure supplement 1E**). This agreed with structural studies where binding between MLL1C and the nucleosome occurs primarily through interactions with DNA and, to a lesser degree, the H4 N-terminal tail (Lee et al., 2021; Park et al., 2019). Thus, the increased H3K4 methylation observed when the H3 N-terminal tail was acetylated is not due to enhanced MLL1C-nucleosome binding. Instead, H3 acetylation likely released the histone tail from the nucleosome, thereby increasing the apparent H3K4 concentration for MLL1C and enhancing methylation.

To definitively explore if the MLL1C methylation of H3K4 in H3tri^ac^ nucleosomes was responding to *cis* and/or *trans* tail acetylation, we synthesized heterotypic substrates with only one H3K4 residue available in the *cis* or *trans* H3ac context (*i.e.*, [H3tri^ac^ • H3K4me3] *vs*. [H3 • H3tetra^ac^]). Using these substrates with MLL1C, we observed >3-fold enhanced methylation (over [H3 • H3] or [H3 • H3K4me3]) when H3 tail acetylation was *cis* but no significant impact when *trans* (**Figure 2D**). This would support a model where H3 acetylation on the same tail releases H3K4 for methylation by MLL1C and appear to exclude a significant contribution from *trans* tail mechanisms.

### Cellular level of H3K4 methylation is coupled to H3 N-terminal tail hyperacetylation

The above data suggested a molecular model for how the H3 N-terminal tail, via *cis* acetylation, becomes available for H3K4 reader binding or enzymatic modification *in vitro*. To determine if such acetylation could function as an accessibility switch *in vivo*, we developed a novel targeted middle-down mass spectrometry method to provide a single molecule quantitative measure of histone tail modification. We applied this method to acid-extracted histones from asynchronous MCF-7 breast cancer cells to measure the relationship between H3K4 methylation and tail acetylation on the same H3 proteoforms (Holt et al., 2021; Smith and Kelleher, 2013). As expected (Garcia et al., 2007; Peach et al., 2012; Young et al., 2009), the absolute amounts of H3K4me3 and higher Kac states (3ac, 4ac, and 5ac) across adjacent lysine residues were extremely low (**Figure 3 - figure supplement 1A**). Nonetheless, H3K4me3 (<1% of total H3) was strictly associated with molecules that contained multiple acetylations (also <1% of total H3; **Figure 3A**). Given this relationship, we next addressed the hypothesis that increased lysine acetylation may release the H3 tail for more effective H3K4 methylation (*i.e*., acetylation precedes methylation). We treated MCF-7 cells with the KDAC inhibitor sodium butyrate and collected samples at multiple timepoints to measure the levels of H3K4 methylation with *cis* acetylation. H3 poly-acetylation rapidly increased upon butyrate treatment (**Figure 3 - figure supplement 1B** and **Supplementary File 2**), as expected (Holt et al., 2019; Young et al., 2009). We observed an increase in all H3K4 methyl states relative to the unmodified state concomitant with increasing states of H3 acetylation (**Figures 3B-C** and **Supplementary File 2**), while H3K4me3 levels most dramatically increased in tandem with the 5ac H3 state (**Figure 3D**). An example of tandem mass spectra at each acetyl degree, showing the C4^+1^ ion series from which K4 stoichiometry is measured, are shown in **Figure 3C**. These findings support a direct link, where acetylation releases nucleosome-bound H3 tails to localized H3K4 methyltransferases for subsequent methylation (**Figure 4**).

**Figure 3.**
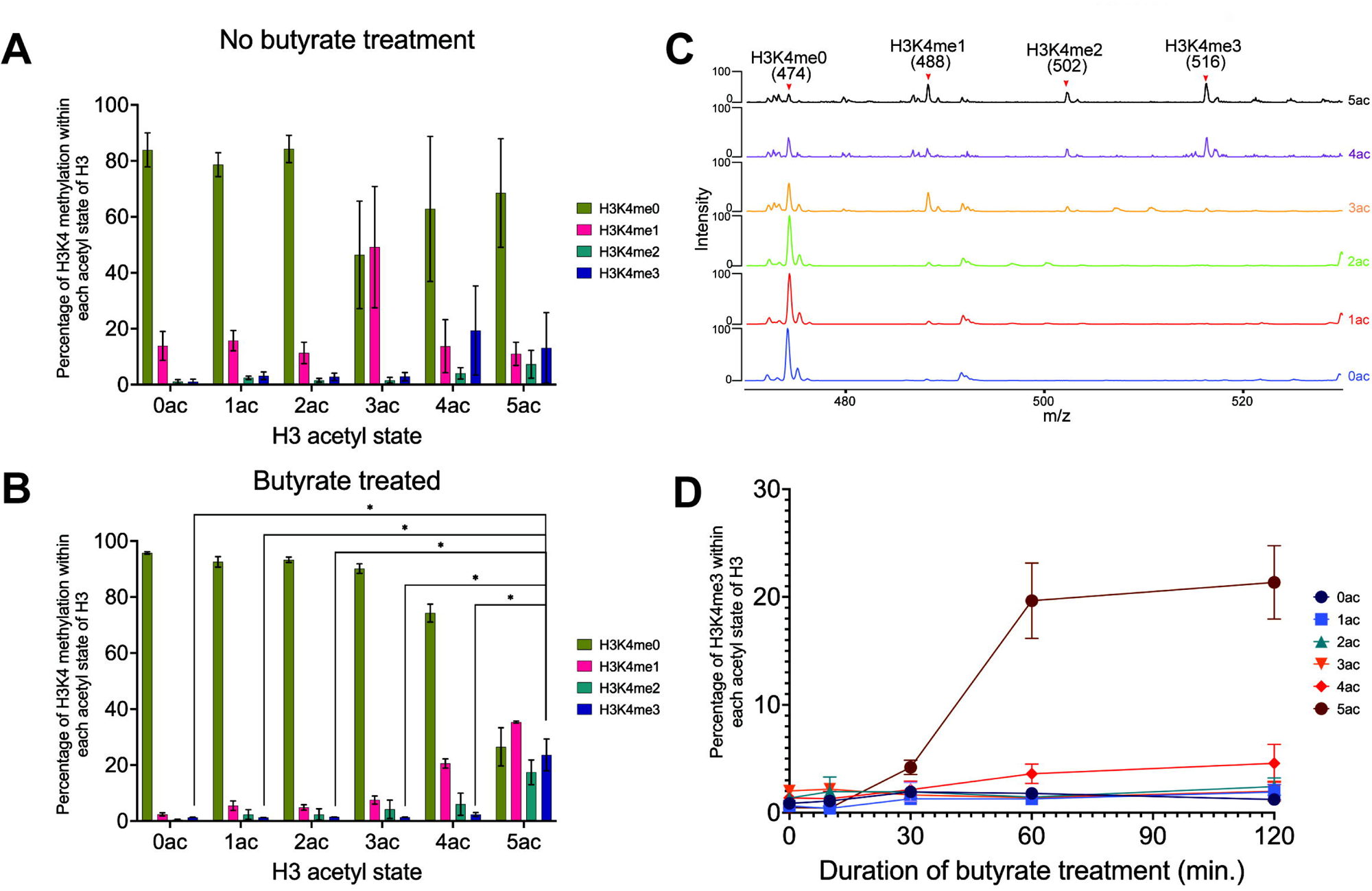
Middle-down MS analysis reveals a hierarchical dependence between H3K4 methylation and cis H3 acetylation in MCF-7 cells. **A)** Occupancy of H3K4 methyl states (me0-1-2-3) within each H3 acetyl state (0ac to 5ac) in asynchronous MCF-7 cells. **B)** Occupancy of H3K4 methyl states within each H3 acetyl state after butyrate treatment (60 mins). Asterisks represent p-values < 0.05. **C)** Representative tandem mass spectra of the targeted C4^+1^ fragment ion series (474 *m/z* unmodified; 488 *m/z* me1; 502 *m/z* me1; 516 *m/z* me3). Each spectrum is an average of MS2 spectra of the indicated H3 acetyl states after 60 minutes butyrate treatment. K4 occupancy stoichiometry is directly correlated with H3 acetylation state and the targeted MS approach provides excellent signal-to-noise for confident quantitation. **D)** Time course of H3K4me3 accumulation with respect to each H3 acetyl state after butyrate treatment. See **Methods** for further information on data acquisition and analysis. All data were collected in biological triplicate with error bars representing S.E.M.

**Figure 4.**
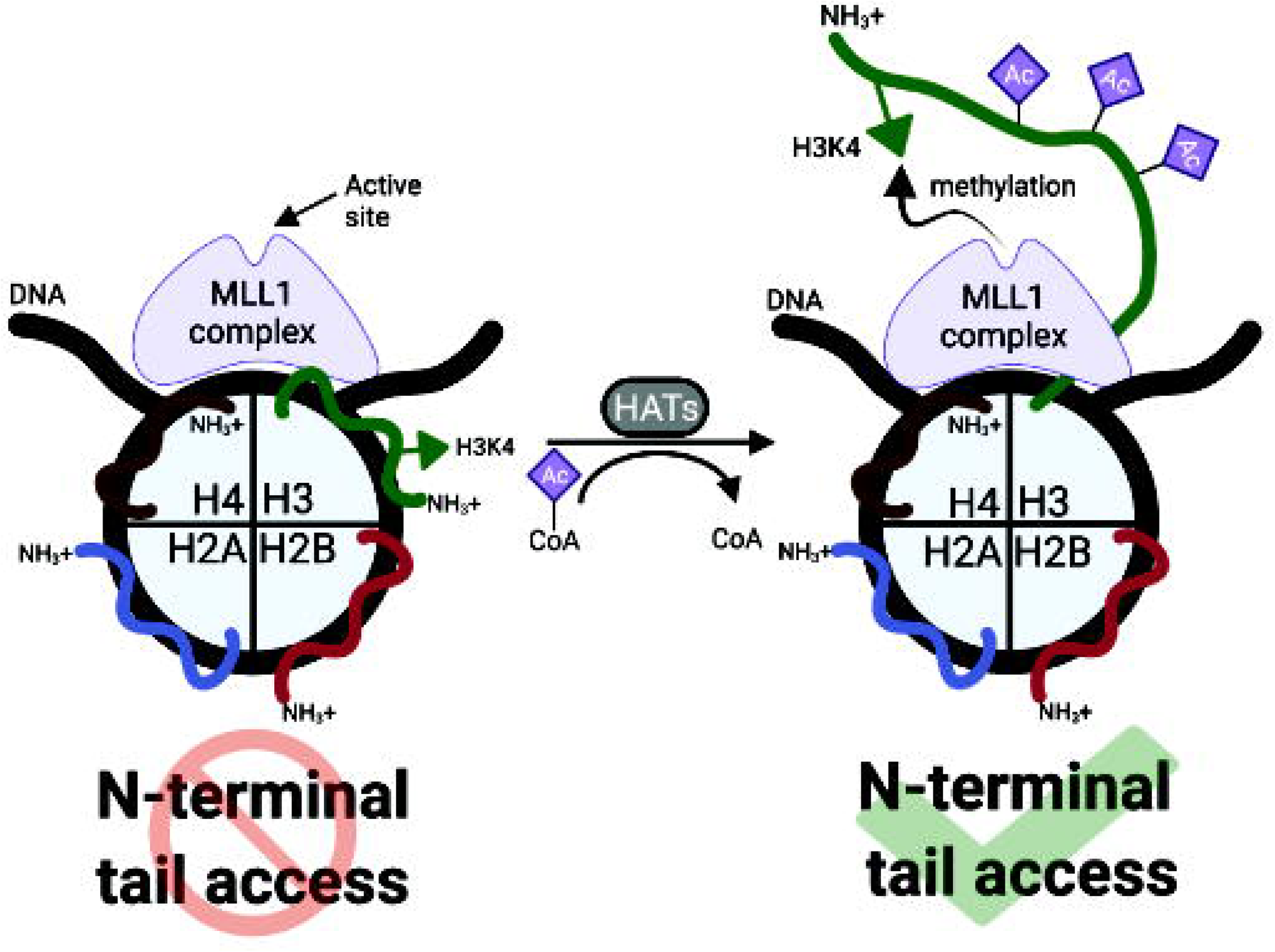
Regulation of H3K4 methylation by *cis*-tail H3 acetylation. Nucleosomal DNA is represented in black; each histone is as labeled, with the core histone N-terminal tails colored to distinguish. MLL1C is in purple.

To explore whether other H3 lysine methyl marks are similarly affected by H3 acetylation, we examined the co-occurrence of H3K9me1-me2-me3 with increasing H3 acetyl states ± butyrate treatment in asynchronous HEK-293 cells (**Figure 3 - figure supplement 1C-D; Supplementary File 3**). In contrast to findings with H3K4me3, H3K9me3 did not accumulate with hyperacetylated H3 proteoforms (*i.e.*, 3ac, 4ac, and 5ac). Recent *in vitro* studies showed tested H3K9 methyltransferases have increased activity to partially acetylated substrates (Trush et al., 2022), so our MS data would suggest this is not a common *in vivo* mechanism: explanations could include that K9 was generally acetylated (and thus occupied) during histone tail release (all penta-acetylated (5ac) H3 proteoforms consisted of acetylation at H3K9, K14, and K18 at minimum (**Figure 3**, **Figure 3 - figure supplement 1E** and **Supplementary File 4**)); or that H3K9 methyltransferases, in contrast to H3K4 methyltransferases, were not generally localized to regions that will accumulate butyrate-induced hyperacetylation.

Taken together, these findings demonstrate a specific *cis*-dependent H3ac regulatory switch that functions to control H3K4 methylation output (**Figure 4**). We posit that such a ‘chromatin-switch’ is important to the ability of cells to translate short-term acetylation signals at gene promoters to longer-term heritable marks of epigenetic memory (Greer et al., 2014; Hörmanseder et al., 2017; Muramoto et al., 2010; Ng et al., 2003).

## DISCUSSION

While previous investigations identified a link between H3K4 methylation and H3 acetylation in diverse species (Garcia et al., 2007; Nightingale et al., 2007; Strahl et al., 1999; Taverna et al., 2007; Young et al., 2009) the molecular basis for this link was unknown, and we posit the H3 acetyl ‘chromatin switch’ defined herein is conserved across eukaryotes. Our new understanding of the dynamic structure of nucleosome histone tails, alternating between collapsed (*i.e.*, nucleosome-bound) and accessible forms (Marunde et al., 2022a; Morrison et al., 2018), has made more plausible the notion that tail availability could be driven by combinatorial *cis* acetylation to directly promote H3K4 methylation. Histone tail lysine acetylation (Kac) can directly recruit residue-specific readers, *e.g.,* bromodomains (Musselman et al., 2012), but acetylation also neutralizes the positive charge on lysine residues and relieves their interaction with negatively charged DNA (*i.e.*, altering the histone tail-DNA binding equilibrium) (Marunde et al., 2022a; Morrison et al., 2018). In the nucleosome context, this decreased histone-DNA binding supports the increased engagement of reader domains that have no direct affinity for the Kac. Conversely, isolated histone tail peptides are ‘constitutively open’ (no DNA to engage), and thus not subject to this mode of regulation.

In this study, we showed that hyperacetylation of the H3 N-terminal tail promoted rapid accumulation of H3K4 methylation in *cis*, most likely by increasing availability of the substrate residue to the MLL1C active site (**Figure 4**). This finding was supported by our *in vitro* enzymatic and *in vivo* mass spectrometric analyses. Methylation assays with MLL1C revealed significantly enhanced enzyme activity towards nucleosome substrates with co-incident acetylated (H3tri^ac^) over unmodified H3 (**Figure 2A-B; Table 1**). dCypher assays demonstrated that the acetylation-mediated increase in H3K4 methylation does not involve stabilized interactions between MLL1C and the nucleosome (**Figure 2D**). And providing major insight, heterotypic nucleosomes containing two distinct PTM-defined forms of histone H3 definitively showed that MLL1C activity was only enhanced in a *cis* H3 N-terminal triacetylation context (**Figure 2D**). In agreement with these *in vitro* findings, middle-down mass spectrometry showed that H3 hyperacetylation and H3K4 methylation co-occurred on the same histone tails in actively cycling cells; furthermore, upon butyrate treatment H3K4 methylation increased for the most highly acetylated proteoforms (**Figure 3**). The *in vivo* relationship required higher degrees of acetylation (preferring at least four acetyl groups per molecule: *e.g.*, H3K9acK14acK18acK23acK27ac) for most effective conversion to the H3K4me3 state. This could be a function of yet to be explained *in vivo* acetylation hierarchies by KATs that are outside the scope of this study but will be important to resolve. Together with binding studies that identify the positive impact of *cis* H3 tail hyperacetylation on H3K4 reader engagement (**Figure 1** and **Figure 1 - figure supplement 1**) (Marunde et al., 2022a; Morrison et al., 2018), our findings suggest a molecular switch that governs when and where the histone H3 N-terminus is available for H3K4-related transactions. Such a switch could be used to establish and heritably maintain the location and function of transcriptional promoters across the genome.

A continued observation from this study is that the binding preference of readers (in this case PHD-fingers that engage histone H3K4) narrows on nucleosomes relative to histone peptides (**Figure 1** and **Figure 1 - figure supplement 1**) (Marunde et al., 2022a; Morgan et al., 2021). Such differences highlight the importance of using a more representative target to identify the most likely/physiologically relevant interactions. However, we also add to the literature confirming the importance of a nucleosome substrate for enzymatic studies (Strelow et al., 2016; Stützer et al., 2016). In this regard our steady-state kinetic methylation assays with MLL1C revealed an intriguing (and previously undescribed) positive cooperativity with its preferred nucleosome substrates (**Figure 2B** *vs*. **Figure 2 - figure supplement 1C**). It is important to consider the suggestion that allosteric factors could regulate interactions between MLL1C and the nucleosome, especially since previous kinetic analyses of this [enzyme: substrate] pair did not address such behavior (Park et al., 2019; Xue et al., 2019). However, positive cooperativity was not evident at the level of [MLL1C: nucleosome] binding, which was independent of substrate identity (*i.e.*, ± H3tri^ac^) at ionic conditions similar to the catalytic assay (**Figure 2C**). Still to be determined are the signals that drive and regulate this cooperativity and its function in MLL1-catalyzed methylation, especially in a higher-order chromatin context.

Taken together, our study highlights a previously unrecognized regulatory mechanism for how writers might engage the histone H3 tail *in vivo*. Although this work focused on H3K4, it will be important to ascertain the consequence of acetylation (or acylation)-mediated changes in accessibility and the function of modifiers and readers of the other lysines on H3, as well as on the other core histones (H2A, H2B, and H4). Underscoring the need for such studies, we note recent *in vitro* analyses employing unmodified nucleosomes sequentially targeted by purified KATs and KMTs suggesting that the acetylation landscape can impact multiple methyltransferases (Trush et al., 2022). While that study reported enhanced G9a activity towards H3K9 as a function of p300-mediated histone acetylation, our mass spectrometric analysis from cells did not show a connection between H3K9me3 and H3 hyperacetylation (**Figure 3 - figure supplement 1C-D**). The field will require a more detailed analysis of the *in vivo* contributions of various KATs/KDACs to regulate histone tail accessibility (in *cis* and/or *trans*) for other chromatin-modifying enzymes to further uncover the molecular details of any sequential histone code.

Finally, we note that multiple studies have identified H3K4 methylation and H3 acetylation as active marks because of their co-occupancy on the promoters and gene bodies of transcribed genes (Santos-Rosa et al., 2002; Strahl et al., 1999; Wozniak and Strahl, 2014). Our findings agree with these observations, but also uncover a previously unrecognized mechanism of H3 cross-talk that impinges on fundamental functions of H3 acetylation and K4 methylation in gene regulation. While this work shows the impact of *cis* tail H3 acetylation on the H3K4 writer MLL1, we note a companion study highlighting how the resulting PTM signature (*cis* >> *trans* H3K4me3tri^ac^) promotes BPTF-nucleosome engagement (Marunde et al., 2022a). Given its central importance, we predict this mechanism may be a target for dysregulation in human disease.

## METHODS

### Expression and purification of GST-tagged PHD reader proteins

Expression constructs for GST-tagged PHD domains from KDM7A (Uniprot #Q6ZMT4; residues 1-100), DIDO1 (Uniprot #Q9BTC0; residues 250-340), and MLL5 (Uniprot #Q8IZD2; residues 100-180) were synthesized in a pGEX-4T-1 vector (*BioMatik Corporation*) and provided by Dr. Mark T. Bedford (UT MD Anderson Cancer Center). Recombinant proteins were expressed and purified as described (Jain et al., 2020). See **Supplementary File 5** for details about the constructs.

### Expression, purification, and assembly of the MLL1 core complex (MLL1C)

Methods for the expression, purification, and assembly of MLL1 core complex (MLL1C: MLL1 SET domain, WDR5, RbBP5, Ash2L and DPY30) were adapted from published protocols (Usher et al., 2021). A polycistronic recombinant expression construct containing the MLL1 SET domain (Uniprot Q03164; residues 3745-3969), WDR5 (Uniprot P61964; residues 2-334), RbBP5 (Uniprot Q15291; residues 1-538) and Ash2L (Uniprot Q9UBL3-3; residues 1-534) in pST44 vector (*aka*. MWRA construct) was a kind gift from Dr. Song Tan (Tan et al., 2005). WDR5 was cloned with an N-terminal hexahistidine (6HIS) tag and TEV protease site to enable purification by immobilized metal affinity chromatography (IMAC), and tag removal by enzymatic cleavage. See **Supplementary File 5** for details about the constructs. Rosetta pLysS *E. coli* cells were transformed with the plasmid and grown on LB plates with 50 μg/mL carbenicillin. Single colonies were used to inoculate 50 mL starter cultures of Terrific Broth (TB) containing 50 μg/mL carbenicillin and grown at 37°C for 16 hours. This culture was transferred to 1L of TB + carbenicillin and grown to OD_600_ ∼0.6 (37°C, 200 RPM, ∼ 4 hours). Recombinant protein expression was induced with 1 mM Isopropyl β-D-1-thiogalactopyranoside (IPTG, *Sigma*; 16°C, 200 RPM, 20 hours). Cells were harvested by centrifugation (5000 RPM, 4°C) and flash frozen in liquid nitrogen.

Frozen cell pellets were resuspended in 50 mL of lysis buffer (50 mM Tris-HCl pH 7.5, 300 mM NaCl, 30 mM imidazole, 3 mM dithiothreitol, and 1 µM ZnCl_2_) containing 0.5 mg/mL lysozyme (*Sigma*), 250 U Pierce Universal Nuclease (*ThermoFisher*), and an EDTA-free protease inhibitor cocktail (*Roche*) and rotated at 4°C for 1 hour. The resultant mixture was then sonicated [five cycles of 30 sec on/30 sec off at 50% output] and centrifuged at 4°C, 15,000 RPM for 35 min. The clarified lysate was flowed over a 5 mL HisTrap nickel column (*Cytiva*) using an AKTA Pure FPLC (*Cytiva*) at 0.5 mL/min; all FPLC steps were conducted at 4°C. Unbound molecules were removed with 20 column volumes of wash buffer (WB: 50 mM Tris-HCl, pH 7.5, 300 mM NaCl, 30 mM imidazole, 3 mM DTT, and 1 µM ZnCl_2_ at 2 mL/min). The 6HIS-tagged MWRA was eluted in a 15-column volume linear gradient from WB to Elution Buffer (WB + 500 mM imidazole) at 2 mL/min. Fractions containing 6HIS-tagged MWRA were identified by SDS-PAGE, pooled, and supplemented with 6HIS-tagged TEV protease (purified as described: Nautiyal and Kuroda, 2018) at a 1:100 enzyme to substrate molar ratio to cleave the 6HIS-tag on WDR5. This mixture was dialyzed against three changes of WB (each 2L for at least four hours at 4°C) and 6HIS-TEV removed from cleaved MWRA via IMAC (Usher et al., 2021). MWRA flow-through protein solution was concentrated to ∼15 mL using a 30 kDa MWCO centrifugal filter (*EMD Millipore*), ensuring not to concentrate to where solution became yellow/cloudy and viscous. MWRA complex was resolved over a HiLoad 16/60 Superdex 200 pg gel filtration (GF) column (*Cytiva*) pre-equilibrated in 20 mM Tris-HCl, pH 7.5, 300 mM NaCl, 1 mM TCEP, and 1 µM ZnCl_2_. Fractions containing stoichiometric MWRA sub-complex were identified by SDS-PAGE, pooled, and concentrated to ∼15 mL.

HisDPY30 was expressed, purified, and cleaved to remove the 6HIS-tag as described (Patel et al., 2009; Usher et al., 2021). A two-fold molar excess of DPY30 was added to the MWRA sub-complex and incubated on ice for 1 hour. Following incubation, the resulting MLL1C was isolated by gel filtration as described for MWRA, with fractions containing the stoichiometric complex identified by SDS-PAGE, pooled, concentrated to ∼10 µM, and flash frozen. For dCypher experiments with MLL1C, the 6HIS-tag was retained on DPY30.

### Peptides

All peptides were synthesized at UNC peptide synthesis core facility (RRID:SCR_017837), using Fmoc solid phase synthesis, on automated peptide synthesizer (PTI Symphony or CEM Liberty Blue). The peptides were purified by preparative RP-HPLC and characterized by MALDI-TOF MS and analytical HPLC.

### PTM-defined nucleosomes

All PTM-defined nucleosomes (**Supplementary file 5**; homotypic unless stated otherwise) were from the dNuc^TM^ or versaNuc^®^ portfolios (*EpiCypher*). PTMs were confirmed by mass-spectrometry and immunoblotting (if an antibody was available) (Goswami et al., 2021; Marunde et al., 2022b, 2022a; Weinberg et al., 2019).

Nucleosomes with methyllysine analogs (MLA) at H3K4 were created by the versaNuc approach (Marunde et al., 2022a). In brief, histone H3 peptides (aa1-31; A29L) containing K4C and any PTMs of interest were site-specifically reacted with the corresponding haloalkylamine: (2-bromoethyl) trimethyl ammonium bromide for KCme3; 2-chloro-N,N-dimethyl-ethylamine hydrochloride for KCme2; or 2-chloroethyl(methyl)ammonium chloride for KCme1) under SN_2_ reaction conditions as previously (Simon, 2010; Simon and Shokat, 2012), and purified for individual ligation to a H3 tailless nucleosome precursor (H3.1NΔ32 assembled on 147bp 5’ biotinylated 601 DNA; #16-0016). The resulting nucleosomes (assembled at 50-100 μg scale) contained minimal free DNA (<5%), undetectable levels of peptide precursor, and ≥90% fully-defined full-length H3.1 (*e.g.,* **Figure 2 – figure supplement 1D-E**).

Heterotypic nucleosomes were assembled from PTM-defined histone octamers containing N-terminally bridged H3 dimers, and the bridge removed in a scarless manner by approaches to be described elsewhere (Manuscript in Preparation). Heterotypic identity was confirmed at all synthesis steps by analyses additional to those used for homotypics, including Nuc-MS on representative final nucleosomes (Schachner et al., 2021). Heterotypic nomenclature describes each PTM-defined histone in the nucleosome, such that [H3K4me3K9acK14acK18ac • H3] *vs*. [H3K4me3 • H3K9acK14acK18ac] contain the same total PTM complement but distributed *cis* or *trans* on the H3 N-termini.

### dCypher assays

*dCypher* binding assays with PTM-defined nucleosomes were performed under standard conditions that titrate query (*e.g.,* epitope-tagged reader domain(s)) to a fixed concentration of target (*e.g.*, biotinylated PTM-defined nucleosome) with the appropriate Alpha Donor and Acceptor beads (*Perkin Elmer*) (Jain et al., 2020; Marunde et al., 2022b; Weinberg et al., 2019). Binding curves [query:target] were generated using a non-linear 4PL curve fit in Prism 9.0 (*GraphPad*). For each query, the relative EC_50_ (EC_50_^rel^) and hillslope values were derived from the best binding target. EC_50_^rel^ is the half maximal signal for the specified target. Where necessary, we excluded query concentration values determined to be beyond a query’s hook point (signal inhibition due to query exceeding bead saturation). The EC_80_^rel^ was selected as the optimal probing concentration for discovery screens because of the robust signal-to-background and to provide the best opportunity to bind targets without saturating the primary target signal. To compute EC_80_^rel^ values, we used the formula EC_F_^rel^ = (F/ (100 – F)^1/H^ x EC_50_^rel^; F = 80 and H = hillslope.

Briefly, 5 μL of GST-tagged reader domain was incubated with 5 μL of 10 nM biotinylated nucleosomes (*e.g., EpiCypher* #16-9001) for 30 minutes at room temperature in 20 mM HEPES pH 7.5, 250 mM NaCl, 0.01% BSA, 0.01% NP-40, 1 mM DTT in a 384-well plate. A mix of 10 uL of 2.5 μg/mL glutathione acceptor beads (*PerkinElmer*, AL109M) and 5 μg/mL streptavidin donor beads (*PerkinElmer*, 6760002) was prepared in 20 mM HEPES pH 7.5, 250 mM NaCl, 0.01% BSA, 0.01% NP-40 and added to each well. The plate was incubated at room temperature in subdued lighting for 60 minutes, and AlphaLISA signal was measured on a PerkinElmer 2104 EnVision (680 nm laser excitation, 570 nm emission filter ± 50 nm bandwidth). Each binding interaction was performed in duplicate in a 20 µL mix in 384 well plates.

MLL1C binding assays (**Figure 2C**) were performed as above except using Nickel-chelate acceptor beads (10 μg/mL; *Perkin Elmer* AL108M), streptavidin donor beads (20 μg/mL; *Perkin Elmer*) and modified assay buffer (20 mM Tris pH 7.5 + 50 mM NaCl, 0.01% BSA, 0.01% NP-40 and 1 mM DTT); [NaCl] was optimized via a titration assay and 50 mM chosen for subsequent analyses.

In vitro *methylation assays*

Methylation assays (Shinsky et al., 2015) were performed at 15°C for three hours using purified MLL1C enzyme and nucleosome substrate in a reaction volume of 20 µL [in 50 mM HEPES, pH 8.0, 1 mM DTT, 1 µM ZnCl_2_; 10 µM of 9:1 *S*-adenosyl-L-methionine (SAM) *p*-toluenesulfonate salt (*Sigma*) to *S*-adenosyl-L-[*methyl*-^3^H]-methionine ([*methyl*-^3^H]-SAM) (*PerkinElmer*)]. Concentrations of MLL1C and NaCl were optimized from 2D-titration methylation assays at [0, 4 and 40 nM MLL1C] and [0, 50 and 300 mM NaCl] with 2 µg of chicken oligo-nucleosome substrate (**Figure 2 – figure supplement 1A-B**) (Morris et al., 2007). It is notable that, across a range of concentrations, MLL1C stability decreases as temperature increases and methyltransferase activity has been reported to be enhanced in sub-physiological NaCl concentrations (Namitz et al., 2019; Shinsky et al., 2015). For endpoint methylation assays, 4 nM MLL1C and 100 nM nucleosome substrates were tested as above. For steady-state kinetics, 4 nM MLL1C was incubated with a nucleosome substrate titration (0, 23.4, 35.1, 52.7, 79, 119, 178, 267 and 400 nM) and reactions quenched with 5 µL of 5X SDS loading dye. For steady-state kinetic assays, 0.5 µg bovine serum albumin was added to each reaction after quenching to act as a loading guide. To analyze methylation, quenched reactions were resolved by 15% Tris-Glycine SDS-PAGE. Gels were stained with Coomassie dye and bands containing mononucleosomes were excised (with serum albumin as a supporting lane marker) and incubated in a solution of 50% Solvable (*PerkinElmer*) and 50% water at 50°C for three hours. Mixture and gel slices were then combined with 10 mL of Hionic-Fluor scintillation fluid (*PerkinElmer*), dark-adapted overnight, and radioactivity measured on a Liquid Scintillation Counter (*Beckman Coulter*).

### Middle-down mass spectrometry of MCF-7 and HEK-293 cells ± KDAC inhibition

MCF-7 breast cancer cells (ATCC HTB-22) were grown in MEM (*Gibco*) supplemented with 10% fetal bovine serum (*VWR*), 100 I.U. penicillin, 100 µg/mL streptomycin (*Corning*), 0.01 mg/mL human recombinant insulin (*Gibco*), and 5 µg/mL plasmocin (*Invivogen*) at 37°C and 5% CO_2_. HEK-293 (ATCC CRL-1573) were grown in DMEM (*Gibco*) supplemented with 10% fetal bovine serum (*Corning*), 100 I.U. penicillin, and 100 µg/mL streptomycin (*Gibco*) at 37°C and 5% CO_2_. Both cell lines were authenticated via STR profiling and confirmed to be *Mycoplasma* negative.

For mass spectrometric (MS) analysis ± KDAC inhibition, cells were cultured in 150 mm dishes to ∼80% confluence and treated with 5 mM sodium butyrate (or equivalent volume of water) in triplicate for 0, 10, 20, 30, 60, 120 mins. Cells were washed with cold PBS (11.9 mM phosphates, 137 mM NaCl, 2.7 mM KCl) to remove residual sodium butyrate, harvested by scraping, and flash frozen in liquid nitrogen. Histones were acid extracted after nuclei isolation as described (Holt et al., 2021). Isolated histones were resuspended in 85 µL 5% acetonitrile, 0.2% trifluoroacetic acid (TFA) and resolved by offline high-performance liquid chromatography (HPLC) as described (Holt et al., 2021). Reverse Phase HPLC fractionation was performed with a U3000 HPLC system (*ThermoFisher*) with a 150 × 2.1–mm Vydac 218TP 3 µm C18 column (*HiChrom* # 218TP3215), at a flowrate of 0.2 mL/min using a linear gradient from 25% B to 60% B in 60 min. The composition of buffers used were A: 5% acetonitrile and 0.2% TFA and B: 95% acetonitrile and 0.188% TFA. After chromatographic separation and fraction collection, histone H3.1 was selected, diluted in 100 mM ammonium acetate (pH = 4) and digested with Glu-C protease (*Roche*) at 10:1 protein:enzyme for 1 h at room temperature prior to mass spectrometric analysis. The digested samples were diluted to 2 µg/µL. Online HPLC was performed on a U3000 RSLC nano Pro-flow system using a C3 column (Zorbax 300SB-C3, 5 µm; *Agilent*). Samples were maintained at 4 °C and 1 µL injected for each analysis using a microliter pickup. A linear 70-minute gradient of 4-15% B was used (Buffer A: 2% acetonitrile, 0.1% formic acid and Buffer B: 98% acetonitrile and 0.1% formic acid) with a flow rate of 0.2 µL/min. The column eluant was introduced into an Orbitrap Fusion Lumos mass spectrometer (*ThermoFisher*) by nano-electrospray ionization. A static spray voltage of 1900 V and an ion transfer tube temperature of 320 °C were set for the source.

A Fusion Lumos mass spectrometer was used to generate MS data. The 9^th^ charge state of histone H3 was targeted for analysis. MS1 analysis was acquired in the orbitrap with a normal mass range and a 60k resolution setting in positive ion mode. An Automatic Gain Control (AGC) target of 5.0E5 with a 200 ms maximum injection time, three microscans, and a scan range of 585-640 *m/z* were used to identify desired ions for fragmentation. MS2 acquisition was performed in both orbitrap and ion trap mode. Both modes used electron transfer dissociation (ETD), a reaction time of 18 ms, and an injection time of 200 ms. A normal scan range was used for the orbitrap mode with a resolution setting of 30k and an AGC target of 5.0E5, with two micro scans. A narrow scan range of 470-530 *m/z*, targeting ions indicative of K4 modification states, was used for the ion trap mode MS2 with an ACG target of 3.0E4, quadrupole isolation, maximum injection time of 100 ms, and eight microscans.

These MS methods were used with two technical replicates per biological replicate (n=3). An MS ion trap mode with a targeted mass list was used to increase sensitivity to identify known low abundance K4me3-containing proteoforms. The ion trap MS2 spectra were averaged for each H3 acetyl state based on known retention times, and the intensities of ions indicative of the K4 methylation states were manually recorded. Retention times for each acetyl state were approximated as 0ac 35-40 min, 1ac 45-55 min, 2ac 55-60 min, 3ac 62-68 min, 4ac 69-73 min, and 5ac 74-78 min. Precursor mass was used as an additional confirmation and filter of correct acetyl state. For scans yielding low signal and high noise (*i.e.,* 5ac at 0 min butyrate treatment), data were manually curated before averaging. Within acetyl states, the relative proportion of fragment ions for unmodified, mono-, di-, and trimethylation of the H3K4 ion at respective *m/z* of 474, 488, 502, and 516 were recorded per MS run. Values were averaged across replicates of the same conditions and normalized to one hundred percent. A two-tailed t-test was used for significance. Raw MS data is available at (ftp://massive.ucsd.edu/MSV000089089/, ftp://massive.ucsd.edu/MSV000091578/).

The MS method used here is highly targeted to most effectively address the mechanistic or single molecule link between H3 acetylation degree and H3K4 occupancy. The strategy used prioritizes optimization of efficient selection of acetyl degree and of the signal-to-noise for the C4^+1^ ion series. This is to the exclusion of other information that can typically be derived from untargeted approaches. For example, because trapping mass spectrometers are limited in the number of ions, we dispose of unnecessary ions to gas phase enrich the C4 ions series. This method provides a direct measure of stoichiometry of K4un, K4me1, K4me2, and K4me3 within an acetyl state.

## DATA ACCESSIBILITY

Raw MS data is publicly available and has been uploaded to the UCSD MassIVE database (ftp://massive.ucsd.edu/MSV000089089/, ftp://massive.ucsd.edu/MSV000091578/). All analyzed data are reported in the manuscript and Supporting Files.

## ACKNOWLEDGEMENTS

KJ is supported by a Postdoctoral Training Fellowship from the National Institutes of Health (NIH; T32CA217824) to the UNC Lineberger Cancer Center and a Postdoctoral Fellowship from the American Cancer Society (PF-20-149-01-DMC). This work was also supported by NIH grants to NLY (R01GM139295, P01AG066606, and R01CA193235), MSC (R01CA140522), *EpiCypher* (R43CA236474, R44GM117683, R44CA214076 and R44GM116584), and to BDS (R35GM126900). We thank colleagues for the generous supply of materials (see **Methods**) and members of the Cosgrove, Young, *EpiCypher,* and Strahl labs for helpful discussions and suggestions.

## AUTHOR CONTRIBUTIONS

KJ, MRM, JMB, MWC, ZWS, NLY, MCK, and BDS conceptualized and managed one or more elements of the study. ZBG, SAH, EFP, MAC, and MJM assembled and validated PTM-defined histones, dNucs and versaNucs. ZBG, HFT, LM and UCO assembled and validated heterotypic dNucs. KK synthesized histone peptides used in homotypic/heterotypic dNuc and versaNucs. MM, JB, SLG, KLR, IKP, NWH, AV, and ENW performed dCypher assays and analyzed data with input from KJ, MCK, and BDS. KEWN and MSC purified MLL1C for enzymatic analyses. KJ and SWC purified his-tagged MLL1C for binding assays. Methylation assays were performed and analyzed by KJ, SWC, and GCF. MCF7 cells were cultured by KJ, and mass spectrometry was performed and analyzed by FJ, KFP, BCT, and NLY. KJ and BDS wrote the first draft of the manuscript and all authors contributed to subsequent editing.

## DECLARATIONS OF INTEREST

MSC owns stock in and serves on the Consultant Advisory Board for Kathera Bioscience Inc. and holds US patents (8,133,690; 8,715,678; and 10,392,423) for compounds/methods for inhibiting SET1/MLL family complexes. *EpiCypher* is a commercial developer and supplier of reagents (*e.g.*, PTM-defined semi-synthetic nucleosomes; dNucs^TM^ and versaNucs^®^) and platforms (*e.g.,* dCypher^®^) used in this study. MCK and BDS are board members of *EpiCypher*. KK owns *EpiCypher* options.

## FIGURE SUPPLEMENT LEGENDS

**Figure 1 - figure supplement 1. dCypher assays with PHD finger reader domains. A)** Binding curves to determine optimal concentration for screening reader <u>queries</u> (GST-KDM7A PHD, GST- DIDO1 PHD, and GST-MLL5 PHD) to indicated PTM-defined nucleosome <u>targets</u>. **B)** Library binding screen with indicated GST-PHD queries and PTM-defined nucleosome or free DNA (147bp (Widom 601 sequence) or 199bp (Widom 601 sequence flanked by 21bp linkers)) targets. Error bars represent the range of two replicates. Key: H3.1N Δ2, H3.1N Δ32 and H4N Δ15 are nucleosomes assembled with histones lacking the indicated N-terminal residues of H3.1 or H4. **C.** Relative EC_50_ (EC_50_^Rel^) binding values (in nM) (calculated as in **Methods**) between GST-PHD reader proteins and select histone peptides; 95% confidence intervals are represented parenthetically. **D.** EC_50_^Rel^ binding values (in nM) between GST-PHD proteins and select nucleosomes; 95% confidence intervals are represented parenthetically. ND = not determined.

**Figure 2 - figure supplement 1. Methylation assays with MLL1C and nucleosome substrate. A)** MLL1C activity on 2 μg chicken oligonucleosomes (2 μg) was 2D-titration tested by enzyme [0, 4 and 40 nM] and salt [0, 50 and 300 mM NaCl] in the presence of [*methyl*-^3^H]-SAM donor. Methylation (in fmol) is graphed as a function of [MLL1C]. N=2 and error bars are S.D. **B)** Replicate SDS-PAGE Coomassie-stained gels for methylation data from panel **A**. Resolved subunits of MLL1C and nucleosomes are indicated. **C)** Steady-state kinetic data for MLL1C activity with nucleosome substrates as shown in Figure 2B but fit to the Michaelis-Menten equation. **D)** Endpoint methylation activity of MLL1C (4 nM) with MLA (methyl-lysine analog) nucleosome substrates (100 nM) in triplicate; error bars are S.D. **E)** Replicate SDS-PAGE Coomassie-stained gels for methylation data from panel **D**; Lanes marked with “*” and “+” in the Replicate 3 gel are switched. **F)** Replicate SDS-PAGE Coomassie-stained gels for methylation data from Figure 2B and Figure 2 **- figure supplement 1C**. 0.5 μg BSA was loaded as a lane marker to each quenched reaction (except for H3K4me3tri^ac^). **G.** SDS-PAGE Coomassie-stained gel for methylation data from Figure 2C. **H.** EC_50_^rel^ binding values (in nM) between MLL1C and select nucleosomes; 95% confidence intervals (CI) are represented parenthetically. Note: gel images in grayscale and blue are both Coomassie-stained gels. All data were plotted using GraphPad Prism 9.0.

**Figure 3 - figure supplement 1. H3 acetylation states with sodium butyrate treatment. A - B)** Each acetyl state of acid-extracted H3 from 0ac to 5ac is represented as a percent of total H3 before / after 60-minute treatment of asynchronous MCF-7 cells with 5 mM sodium butyrate. **C.** Occupancy of H3K4 methyl states (me0-1-2-3) within each H3 acetyl state (0ac to 5ac) in asynchronous HEK-293 cells. **D.** Occupancy of H3K4 methyl states within each H3 acetyl state after butyrate treatment (120 mins) in asynchronous HEK-293 cells. **E.** Percentage of {K9ac, K14ac, K18ac} proteoforms within 3-5ac acetyl states of H3.1 ± butyrate treatment in HEK-293 cells. N=3 and error bars are S.E.M. for all data.

### SUPPLEMENTARY FILE LEGENDS

**Supplementary File 1.** Steady-state Michaelis-Menten kinetic parameters ^a^ = methylation signal observed was indistinguishable from background and thus kinetic parameters could not be determined.

**Supplementary File 2.** Mass Spectrometric data for H3K4 methylation and H3 acetylation in asynchronous MCF-7 cells as a function of butyrate treatment. (See Excel file).

**Supplementary File 3.** Mass Spectrometric data for H3K9 methylation and H3 acetylation in asynchronous HEK-293 cells as a function of butyrate treatment. (See Excel file).

**Supplementary File 4.** Mass Spectrometric data for H3K9acK14acK18ac abundance within H3 acetyl states in asynchronous HEK-293 cells as a function of butyrate treatment. (See Excel file).

**Supplementary File 5.** A table resources detailing recombinant protein constructs, peptides, and nucleosomes used in this study. (See Excel file).

## Notes

ftp://massive.ucsd.edu/MSV000089089/

ftp://massive.ucsd.edu/MSV000091578/

### Summary of Updates

In vitro methylation data with heterotypic nucleosomes added (Figure 2D). Additional H3K9me and H3K9ac mass spec data included (Figure 3- figure supplement 1). Author list updated.

## REFERENCES

Garcia BA, Pesavento JJ, Mizzen CA, Kelleher NL. 2007. Pervasive combinatorial modification of histone H3 in human cells. Nat Methods 4:487–489. doi:10.1038/nmeth1052

Ghoneim M, Fuchs HA, Musselman CA. 2021. Histone Tail Conformations: A Fuzzy Affair with DNA. Trends Biochem Sci. doi:10.1016/j.tibs.2020.12.012

Goswami J, MacArthur T, Bailey K, Spears G, Kozar RA, Auton M, Dong JF, Key NS, Heller S, Loomis E, Hall NW, Johnstone AL, Park MS. 2021. Neutrophil Extracellular Trap Formation and Syndecan-1 Shedding Are Increased After Trauma. Shock 56:433–439. doi:10.1097/SHK.0000000000001741

Greer EL, Beese-Sims SE, Brookes E, Spadafora R, Zhu Y, Rothbart SB, Aristizábal-Corrales D, Chen S, Badeaux AI, Jin Q, Wang W, Strahl BD, Colaiácovo MP, Shi Y. 2014. A Histone Methylation Network Regulates Transgenerational Epigenetic Memory in C. elegans. Cell Rep 7:113–126. doi:10.1016/J.CELREP.2014.02.044

Holt M V, Wang T, Young N. 2021. Expeditious Extraction of Histones from Limited Cells or Tissue Samples and Quantitative Top-Down Proteomic Analysis. Curr Protoc 1. doi:10.1002/CPZ1.26

Holt M V., Wang T, Young NL. 2019. High-Throughput Quantitative Top-Down Proteomics: Histone H4. J Am Soc Mass Spectrom 30:2548–2560. doi:10.1007/s13361-019-02350-z

Hörmanseder E, Simeone A, Allen GE, Bradshaw CR, Figlmüller M, Gurdon J, Jullien J. 2017. H3K4 Methylation-Dependent Memory of Somatic Cell Identity Inhibits Reprogramming and Development of Nuclear Transfer Embryos. Cell Stem Cell 21:135–143.e6. doi:10.1016/J.STEM.2017.03.003

Jain K, Fraser CS, Marunde MR, Parker MM, Sagum C, Burg JM, Hall N, Popova IK, Rodriguez KL, Vaidya A, Krajewski K, Keogh M-C, Bedford MT, Strahl BD. 2020. Characterization of the plant homeodomain (PHD) reader family for their histone tail interactions. Epigenetics & Chromatin 2020 13:1 13:1–11. doi:10.1186/S13072-020-0328-Z

Jenuwein T, Allis CD. 2001. Translating the Histone Code. Science (1979) 293:1074–1080. doi:10.1126/science.1063127

Lee YT, Ayoub A, Park SH, Sha L, Xu J, Mao F, Zheng W, Zhang Y, Cho US, Dou Y. 2021. Mechanism for DPY30 and ASH2L intrinsically disordered regions to modulate the MLL/SET1 activity on chromatin. Nat Commun 12. doi:10.1038/s41467-021-23268-9

Marunde MR, Fuchs HA, Burg JM, Popova IK, Vaidya A, Hall NW, Meiners MJ, Watson R, Howard SA, Novitzky K, McAnarney E, Cheek MA, Sun Z-W, Venters BJ, Keogh M-C, Musselman CA. 2022a. Nucleosome conformation dictates the histone code. bioRxiv 2022.02.21.481373. doi:10.1101/2022.02.21.481373

Marunde MR, Popova IK, Weinzapfel EN, Keogh M-C. 2022b. The dCypher Approach to Interrogate Chromatin Reader Activity Against Posttranslational Modification-Defined Histone Peptides and Nucleosomes 231–255. doi:10.1007/978-1-0716-2140-0_13

Morgan MAJ, Popova IK, Vaidya A, Burg JM, Marunde MR, Rendleman EJ, Dumar ZJ, Watson R, Meiners MJ, Howard SA, Khalatyan N, Vaughan RM, Rothbart SB, Keogh M-C, Shilatifard A. 2021. A trivalent nucleosome interaction by PHIP/BRWD2 is disrupted in neurodevelopmental disorders and cancer. Genes Dev. doi:10.1101/GAD.348766.121

Morris SA, Rao B, Garcia BA, Hake SB, Diaz RL, Shabanowitz J, Hunt DF, Allis CD, Lieb JD, Strahl BD. 2007. Identification of histone H3 lysine 36 acetylation as a highly conserved histone modification. Journal of Biological Chemistry 282:7632–7640. doi:10.1074/jbc.M607909200

Morrison EA, Bowerman S, Sylvers KL, Wereszczynski J, Musselman CA. 2018. The conformation of the histone H3 tail inhibits association of the BPTF PHD finger with the nucleosome. Elife 7. doi:10.7554/eLife.31481

Muramoto T, Müller I, Thomas G, Melvin A, Chubb JR. 2010. Methylation of H3K4 Is Required for Inheritance of Active Transcriptional States. Current Biology 20:397–406. doi:10.1016/J.CUB.2010.01.017

Musselman CA, Lalonde M-E, Côté J, Kutateladze TG. 2012. Perceiving the epigenetic landscape through histone readers. Nature Structural & Molecular Biology 2012 19:12 19:1218–1227. doi:10.1038/nsmb.2436

Namitz KEW, Tan S, Cosgrove MS. 2019. Hierarchical assembly of the MLL1 core complex within a biomolecular condensate regulates H3K4 methylation. bioRxiv. doi:10.1101/870667

Nautiyal K, Kuroda Y. 2018. A SEP tag enhances the expression, solubility and yield of recombinant TEV protease without altering its activity. N Biotechnol 42:77–84. doi:10.1016/j.nbt.2018.02.006

Ng HH, Robert F, Young RA, Struhl K. 2003. Targeted Recruitment of Set1 Histone Methylase by Elongating Pol II Provides a Localized Mark and Memory of Recent Transcriptional Activity. Mol Cell 11:709–719. doi:10.1016/S1097-2765(03)00092-3

Nightingale KP, Gendreizig S, White DA, Bradbury C, Hollfelder F, Turner BM. 2007. Cross-talk between Histone Modifications in Response to Histone Deacetylase Inhibitors: MLL4 LINKS HISTONE H3 ACETYLATION AND HISTONE H3K4 METHYLATION. Journal of Biological Chemistry 282:4408–4416. doi:10.1074/JBC.M606773200

Park SH, Ayoub A, Lee YT, Xu J, Kim H, Zheng W, Zhang B, Sha L, An S, Zhang Y, Cianfrocco MA, Su M, Dou Y, Cho US. 2019. Cryo-EM structure of the human MLL1 core complex bound to the nucleosome. Nat Commun 10. doi:10.1038/s41467-019-13550-2

Patel A, Dharmarajan V, Vought VE, Cosgrove MS. 2009. On the mechanism of multiple lysine methylation by the human mixed lineage leukemia protein-1 (MLL1) core complex. Journal of Biological Chemistry 284:24242–24256. doi:10.1074/jbc.M109.014498

Patel A, Vought VE, Dharmarajan V, Cosgrove MS. 2011. A Novel Non-SET Domain Multi-subunit Methyltransferase Required for Sequential Nucleosomal Histone H3 Methylation by the Mixed Lineage Leukemia Protein-1 (MLL1) Core Complex *. Journal of Biological Chemistry 286:3359–3369. doi:10.1074/jbc.M110.174524

Peach SE, Rudomin EL, Udeshi ND, Carr SA, Jaffe JD. 2012. Quantitative Assessment of Chromatin Immunoprecipitation Grade Antibodies Directed against Histone Modifications Reveals Patterns of Co-occurring Marks on Histone Protein Molecules. Molecular & Cellular Proteomics 11:128–137. doi:10.1074/MCP.M111.015941

Rao RC, Dou Y. 2015. Hijacked in cancer: The KMT2 (MLL) family of methyltransferases. Nat Rev Cancer. doi:10.1038/nrc3929

Rossetto D, Avvakumov N, Côté J. 2012. Histone phosphorylation. Epigenetics 7:1098–1108. doi:10.4161/epi.21975

Santos-Rosa H, Schneider R, Bannister AJ, Sherriff J, Bernstein BE, Emre NCT, Schreiber SL, Mellor J, Kouzarides T. 2002. Active genes are tri-methylated at K4 of histone H3. Nature 2002 419:6905 419:407–411. doi:10.1038/nature01080

Schachner LF, Jooß K, Morgan MA, Piunti A, Meiners MJ, Kafader JO, Lee AS, Iwanaszko M, Cheek MA, Burg JM, Howard SA, Keogh MC, Shilatifard A, Kelleher NL. 2021. Decoding the protein composition of whole nucleosomes with Nuc-MS. Nature Methods 2021 18:3 18:303–308. doi:10.1038/s41592-020-01052-9

Sha L, Ayoub A, Cho US, Dou Y. 2020. Insights on the regulation of the MLL/SET1 family histone methyltransferases. Biochim Biophys Acta Gene Regul Mech. doi:10.1016/j.bbagrm.2020.194561

Shinsky SA, Monteith KE, Viggiano S, Cosgrove MS. 2015. Biochemical reconstitution and phylogenetic comparison of human SET1 family core complexes involved in histone methylation. Journal of Biological Chemistry 290:6361–6375. doi:10.1074/jbc.M114.627646

Simon MD. 2010. Installation of Site-Specific Methylation into Histones Using Methyl Lysine Analogs. Curr Protoc Mol Biol 90:21.18.1–21.18.10. doi:10.1002/0471142727.MB2118S90

Simon MD, Shokat KM. 2012. A Method to Site-Specifically Incorporate Methyl-Lysine Analogues into Recombinant Proteins. Methods Enzymol 512:57–69. doi:10.1016/B978-0-12-391940-3.00003-2

Smith LM, Kelleher NL. 2013. Proteoform: a single term describing protein complexity. Nature Methods 2013 10:3 10:186–187. doi:10.1038/nmeth.2369

Strahl BD, Allis CD. 2000. The language of covalent histone modifications. Nature 403:41–45. doi:10.1038/47412

Strahl BD, Ohba R, Cook RG, Allis CD. 1999. Methylation of histone H3 at lysine 4 is highly conserved and correlates with transcriptionally active nuclei in Tetrahymena. Proceedings of the National Academy of Sciences 96:14967–14972. doi:10.1073/PNAS.96.26.14967

Strelow JM, Xiao M, Cavitt RN, Fite NC, Margolis BJ, Park KJ. 2016. The Use of Nucleosome Substrates Improves Binding of SAM Analogs to SETD8. J Biomol Screen 21:786–794. doi:10.1177/1087057116656596

Stützer A, Liokatis S, Kiesel A, Schwarzer D, Sprangers R, Söding J, Selenko P, Fischle W. 2016. Modulations of DNA Contacts by Linker Histones and Post-translational Modifications Determine the Mobility and Modifiability of Nucleosomal H3 Tails. Mol Cell 61:247–259. doi:10.1016/J.MOLCEL.2015.12.015/ATTACHMENT/48EB5ADD-404B-4984-923B-7C6FC2E26120/MMC1.PDF

Su Z, Denu JM. 2016. Reading the Combinatorial Histone Language. ACS Chem Biol 11:564–574. doi:10.1021/acschembio.5b00864

Tan S, Kern RC, Selleck W. 2005. The pST44 polycistronic expression system for producing protein complexes in Escherichia coli. Protein Expr Purif 40:385–395. doi:10.1016/j.pep.2004.12.002

Taverna SD, Ueberheide BM, Liu Y, Tackett AJ, Dias RL, Shabanowitz J, Chait BT, Hunt DF, Allis CD. 2007. Long-distance combinatorial linkage between methylation and acetylation on histone H3 N termini. Proc Natl Acad Sci U S A 104:2086–2091. doi:10.1073/pnas.0610993104

Taylor BC, Young NL. 2021. Combinations of histone post-translational modifications. Biochemical Journal 478:511–532. doi:10.1042/BCJ20200170

Trush V V., Feller C, Li ASM, Allali-Hassani A, Szewczyk MM, Chau I, Eram MS, Jiang B, Luu R, Zhang F, Barsyte-Lovejoy D, Aebersold R, Arrowsmith CH, Vedadi M. 2022. Enzymatic nucleosome acetylation selectively affects activity of histone methyltransferases in vitro. Biochimica et Biophysica Acta (BBA) - Gene Regulatory Mechanisms 1865:194845. doi:10.1016/J.BBAGRM.2022.194845

Usher ET, Namitz KEW, Cosgrove MS, Showalter SA. 2021. Probing multiple enzymatic methylation events in real time with NMR spectroscopy. Biophys J 120:4710–4721. doi:10.1016/J.BPJ.2021.09.034

Vaughan RM, Kupai A, Rothbart SB. 2021. Chromatin Regulation through Ubiquitin and Ubiquitin-like Histone Modifications. Trends Biochem Sci. doi:10.1016/j.tibs.2020.11.005

Weinberg DN, Papillon-Cavanagh S, Chen H, Yue Y, Chen X, Rajagopalan KN, Horth C, McGuire JT, Xu X, Nikbakht H, Lemiesz AE, Marchione DM, Marunde MR, Meiners MJ, Cheek MA, Keogh M-C, Bareke E, Djedid A, Harutyunyan AS, Jabado N, Garcia BA, Li H, Allis CD, Majewski J, Lu C. 2019. The histone mark H3K36me2 recruits DNMT3A and shapes the intergenic DNA methylation landscape. Nature 573:281–286. doi:10.1038/s41586-019-1534-3

Weinberg DN, Rosenbaum P, Chen X, Barrows D, Horth C, Marunde MR, Popova IK, Gillespie ZB, Keogh MC, Lu C, Majewski J, Allis CD. 2021. Two competing mechanisms of DNMT3A recruitment regulate the dynamics of de novo DNA methylation at PRC1-targeted CpG islands. Nat Genet 53:794–800. doi:10.1038/s41588-021-00856-5

Weiss JN. 1997. The Hill equation revisited: uses and misuses. The FASEB journal: official publication of the Federation of American Societies for Experimental Biology 11:835–841. doi:0892-6638/97/0011

Wozniak G, Strahl B. 2014. Hitting the “mark”: interpreting lysine methylation in the context of active transcription. Biochim Biophys Acta 1839:1353–1361. doi:10.1016/J.BBAGRM.2014.03.002

Xue H, Yao T, Cao M, Zhu G, Li Y, Yuan G, Chen Y, Lei M, Huang J. 2019. Structural basis of nucleosome recognition and modification by MLL methyltransferases. Nature 573:445–449. doi:10.1038/s41586-019-1528-1

Young NL, DiMaggio PA, Garcia BA. 2010. The significance, development and progress of high-throughput combinatorial histone code analysis. Cellular and Molecular Life Sciences 2010 67:23 67:3983–4000. doi:10.1007/S00018-010-0475-7

Young NL, DiMaggio PA, Plazas-Mayorca MD, Baliban RC, Floudas CA, Garcia BA. 2009. High throughput characterization of combinatorial histone codes. Molecular and Cellular Proteomics 8:2266–2284. doi:10.1074/mcp.M900238-MCP200

