## Supplementary figures and images for "An acetylation-mediated chromatin switch governs H3K4 methylation read-write capability"

### Figure 1-Figure Supplement 1

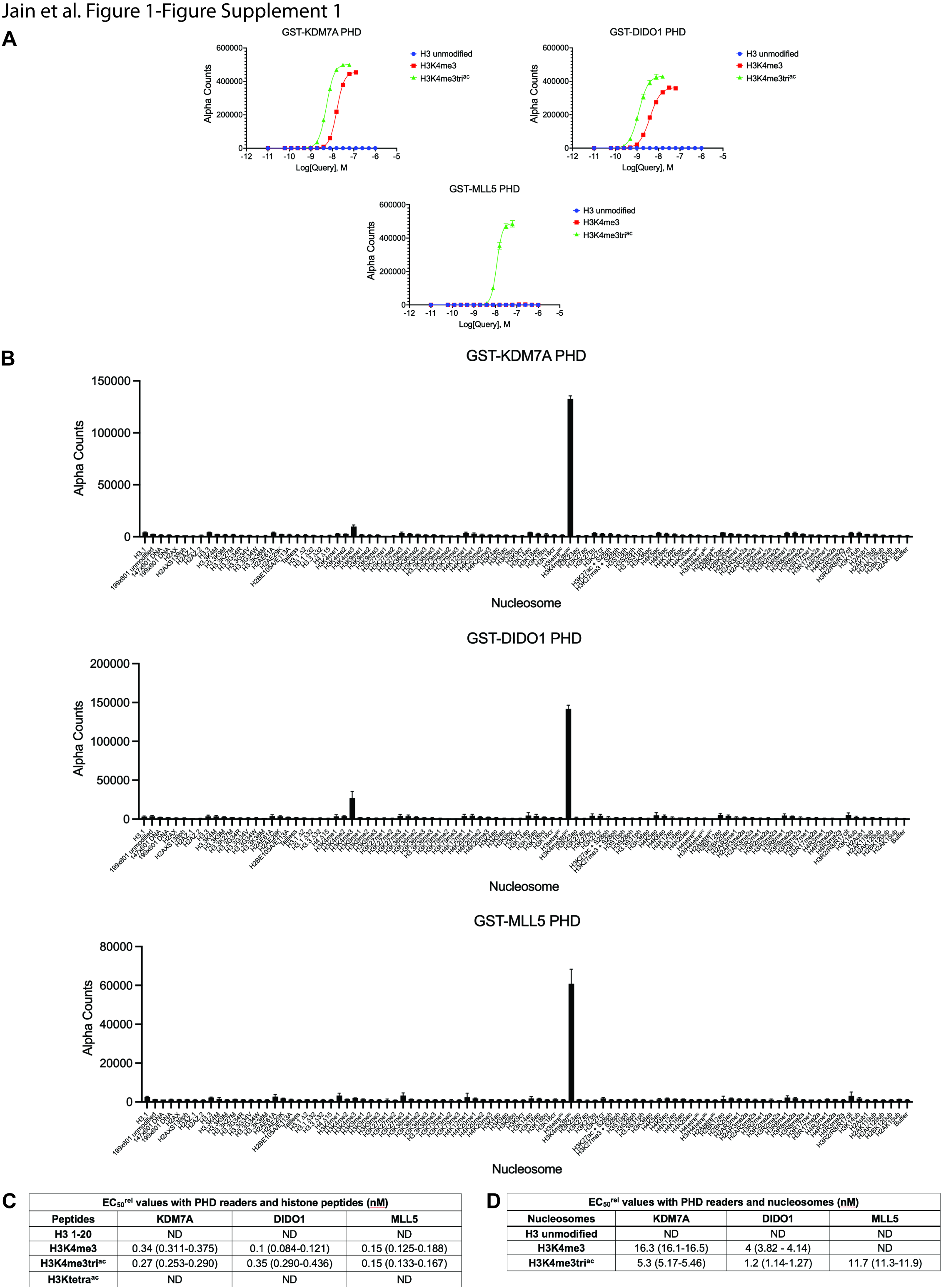

### Figure 2-Figure Supplement 1

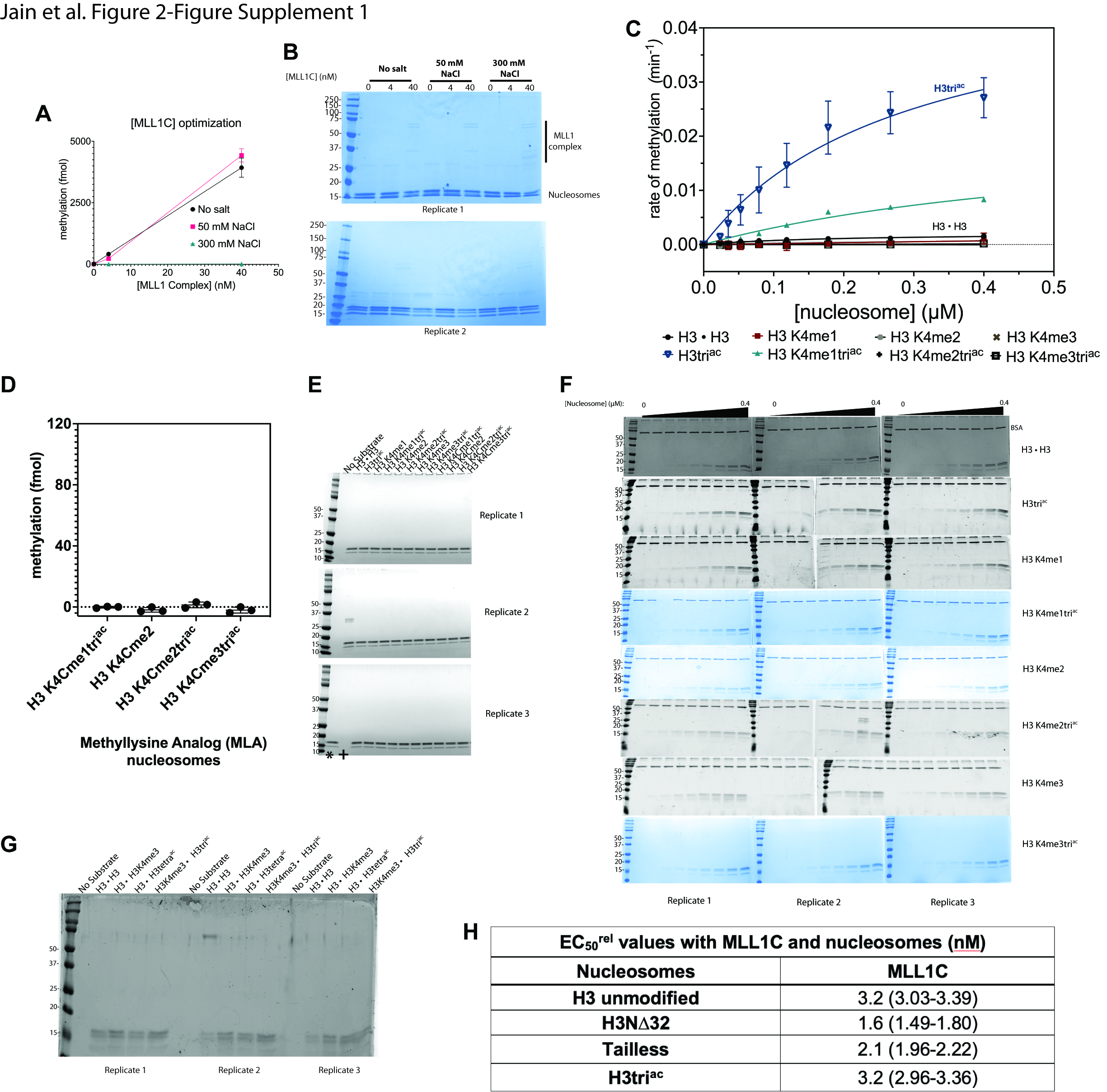

### Figure 3-Figure Supplement 1

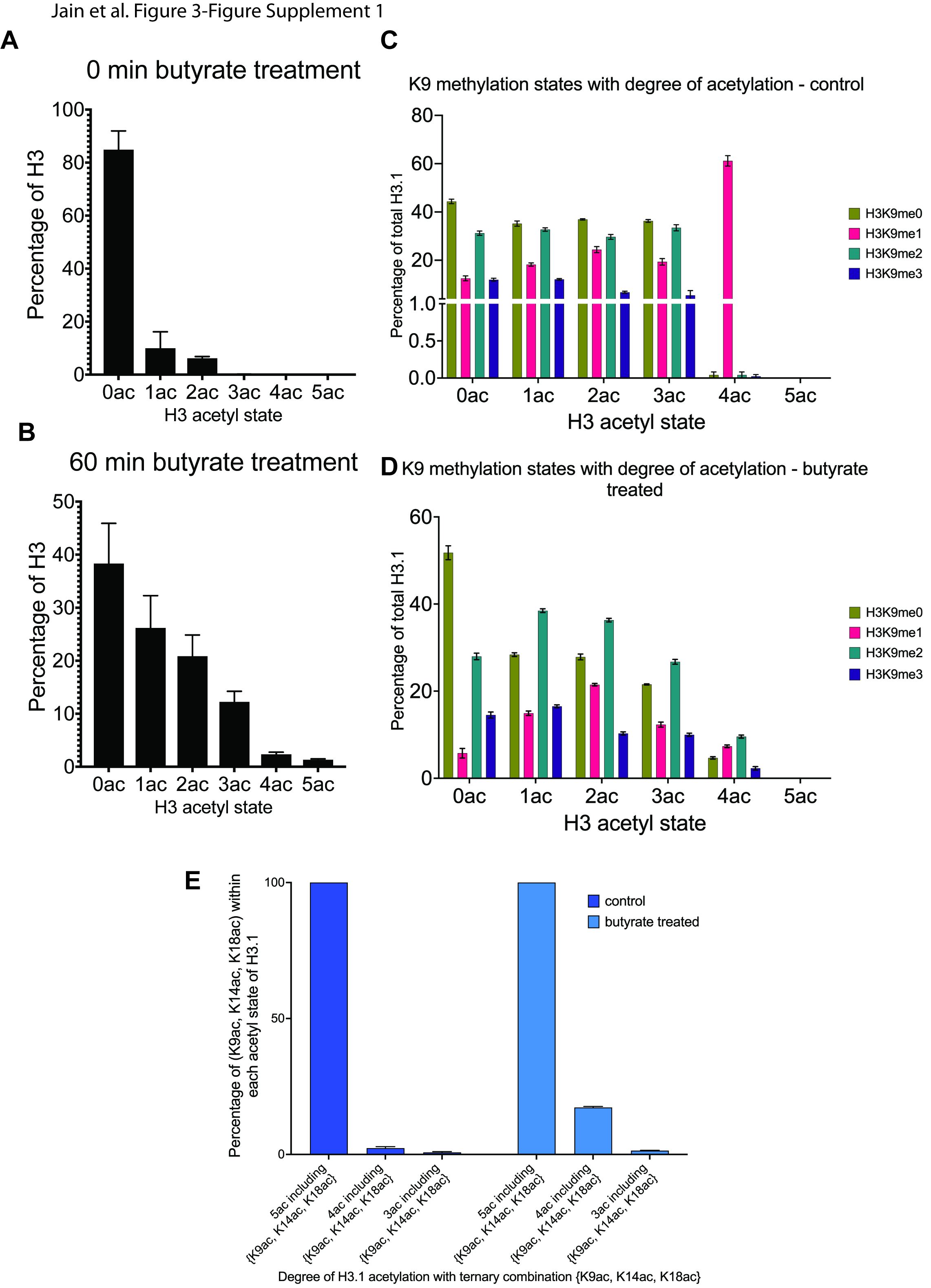
